# On the decidability of population size histories from finite allele frequency spectra

**DOI:** 10.1101/235317

**Authors:** Soheil Baharian, Simon Gravel

## Abstract

Understanding the historical events that shaped current genomic diversity has applications in historical, biological, and medical research. However, the amount of historical information that can be inferred from genetic data is finite, which leads to an identifiability problem. For example, different historical processes can lead to identical distribution of allele frequencies. This identifiability issue casts a shadow of uncertainty over the results of any study which uses the frequency spectrum to infer past demography. It has been argued that imposing mild ‘reasonableness’ constraints on demographic histories can enable unique reconstruction, at least in an idealized setting where the length of the genome is nearly infinite. Here, we discuss this problem for finite sample size and genome length. Using the diffusion approximation, we obtain bounds on likelihood differences between similar demographic histories, and use them to construct pairs of very different reasonable histories that produce almost-identical frequency distributions. The finite-genome problem therefore remains poorly determined even among reasonable histories. Where fits to few-parameter models produce narrow parameter confidence intervals, large uncertainties lurk hidden by model assumption.

## 1 Introduction

Genetic variation across individuals contains information about the evolutionary and demographic history of populations. A simple and efficient summary statistic of genomic variation commonly used in inference studies of population demography is the allele frequency spectrum, describing the proportion of segregating sites as a function of the population frequency of the derived allele [1, 2, 3, 4, 5, 6]. Several computational models have been proposed to reconstruct historical population sizes that are consistent with observed allele frequency spectra [7, 8, 9, 10, 11, 12, 13]. Under the assumption of a neutral Wright-Fisher model, these inferred histories are often taken to be representative of the effective historical population sizes [14, 15, 16, 11, 17]. They are also used as baseline models to identify regions under selection [18, 19, 20] and to predict patterns of deleterious variation in human genomes [21, 22, 23].

However, Myers *et al.* [24] showed that the solution to this inference problem is not unique. To illustrate this, they constructed a family of distinct demographic histories whose frequency spectra under neutral Wright-Fisher evolution are identical for any sample size. This poses a serious practical challenge, since a demographic model that fits well the observed neutral diversity is not guaranteed to be historically accurate or to provide an appropriate model for deleterious variation.

On the other hand, Bhaskar and Song [25] have argued that the families of demographic models constructed by Myers *et al.* are not biologically realistic, because they require historical population sizes that oscillate on arbitrarily short time-scales. They proved that we can uniquely reconstruct the underlying demography from the allele frequency spectrum if (i) we limit our search to historical population sizes that are piecewise-continuous functions of time with a given maximum number of oscillations, (ii) we have enough samples, and (iii) we assume an infinitely long genome [25].

Refs [24, 25] are both correct, but they send different messages regarding the reliability of inferred histories. Does the problem of identifiability raised by [24] bear on applied inference, or should it be considered a purely theoretical result about a class of pathological functions with little biological relevance?

In this article, we seek to resolve this question by addressing the identifiability problem in more realistic scenarios where both sample size and genome length are finite. Recent work by Terhorst and Song [6] has started to address this question by providing strict bounds on the accuracy of demographic inference based on the allele frequency spectrum. In particular, they have focused on the possibility of reconstructing history prior to a bottleneck, which is challenging because of the lost diversity (and thus lost information) during the bottleneck.

Extending this work to situations with arbitrary demographies, we argue that the problem remains poorly determined, even without bottlenecks, in the sense that vastly different population histories can produce statistically indistinguishable allele frequency spectra. Ancient history differences are most difficult to detect, as expected, but we also explain how the approach of Myers *et al.* can be modified to construct well-behaved, practically indistinguishable histories with somewhat more recent differences.

Our arguments are based on two simple observations. First, similar histories should produce similar frequency spectra. Second, the Myers *et al.* family of functions may exhibit infinitely fast oscillations, but, given the extremely small amplitude of these oscillations, they can be replaced by smooth, non-oscillating functions with tiny effect on the frequency spectrum. Thus macroscopically different histories can produce microscopic differences in the frequency distribution. These small differences could in principle be detected given an infinitely long genome, as per the Bhaskar and Song result, but they could not be detected given a finite genome of realistic length. To prove this, we first produce upper bounds on the differences between frequency spectra produced by two similar demographic models, and on the likelihood ratio between the two models given an observed frequency spectrum. Using these bounds and the family of functions given by Myers *et al.,* we construct a family of plausible demographic histories that are very distinct but practically indistinguishable.

The main practical message from this study is that any demographic inference study based on the frequency spectrum *must* have large zones of uncertainty. These can be detected, in principle, by exploring the likelihood surface over the space of all possible functions. However, most inference studies use few-parameter demographic models and estimate parameter uncertainties assuming that these models are correct. In such studies, an excellent fit with small parameter uncertainties can still mask a model that is completely wrong.

This paper is organized as follows. We present an intuitive discussion regarding identifiable demographies in Sec. 2, discussing the properties of the construction of Myers *et al.* [24]. In Sec. 3, we provide the preliminary theory necessary for our analysis. We formally derive a bound on the change in the allele frequency spectrum due to a change in the population size history in Sec. 4, both for infinite and finite genome length, and discuss the relationship with Terhorst and Song [6].

## 2 The diffusion approximation and the identifiability problem

The evolution of the allele frequency spectrum *P*(*y, t*) for a large randomly mating population with Wright-Fisher reproduction can be modeled through a diffusion process [26], which we write as

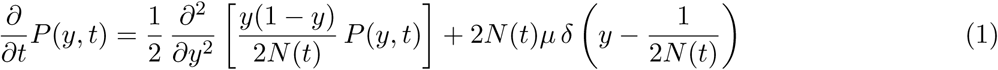

where 0 < *y* < 1 is the allele frequency in the population, *μ* is the mutation rate per generation, *N*(*t*) is the size of the population at time *t* (measured in generations), and *δ*(·) is the Dirac delta function. We assume that the population size is large enough that the frequency *y* can be approximated by a continuous variable. In this formulation, the first term on the right-hand side describes the effect of genetic drift and the second describes that of new mutations entering the population with initial frequency 1/2*N*(*t*). The diffusion equation can also be written in “genetic” or “diffusion” time, 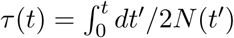, in which drift occurs at a constant rate, i.e.,

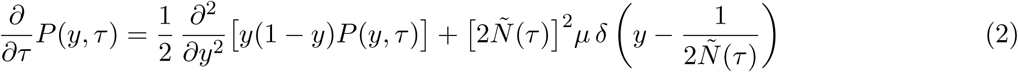

where *Ñ*(*τ*) = *N*(*t*(*τ*)).

The frequency spectrum *P*(*y,τ*) depends on *Ñ*(*τ*) and therefore contains information about the population size history. The problem of identifiability of demographic histories, raised by [24], is that two different histories *Ñ*_1_(*τ*) and *Ñ*_2_(*τ*) can lead to identical frequency spectra. Concretely, Myers *et al.* [24] considered functions *Ñ*_2_(*τ*) of the form

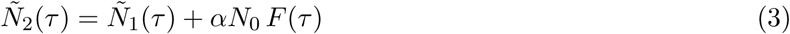

where *α* is a constant and *N*_0_ denotes the current population size. They showed that histories *Ñ*_1_(*τ*) and *Ñ*_2_(*τ*) would lead to identical allele frequency spectra if the function *F*(*τ*) obeys

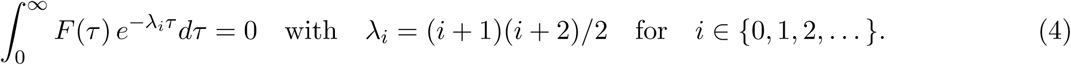

They also showed that such functions exist and constructed an example, 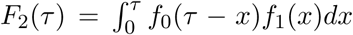 where *f*_0_(*τ*) = exp(−1/*τ*^2^) and 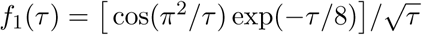, which is displayed in the left panel of Fig. 1 (they set *α* = −9 to ensure that *Ñ*_2_(*τ*) is strictly positive). Since *Ñ*_2_(*τ*) cannot be distinguished from *Ñ*_1_(*τ*) based on the allele frequency spectrum, the inference problem is poorly determined.

**Figure 1:**
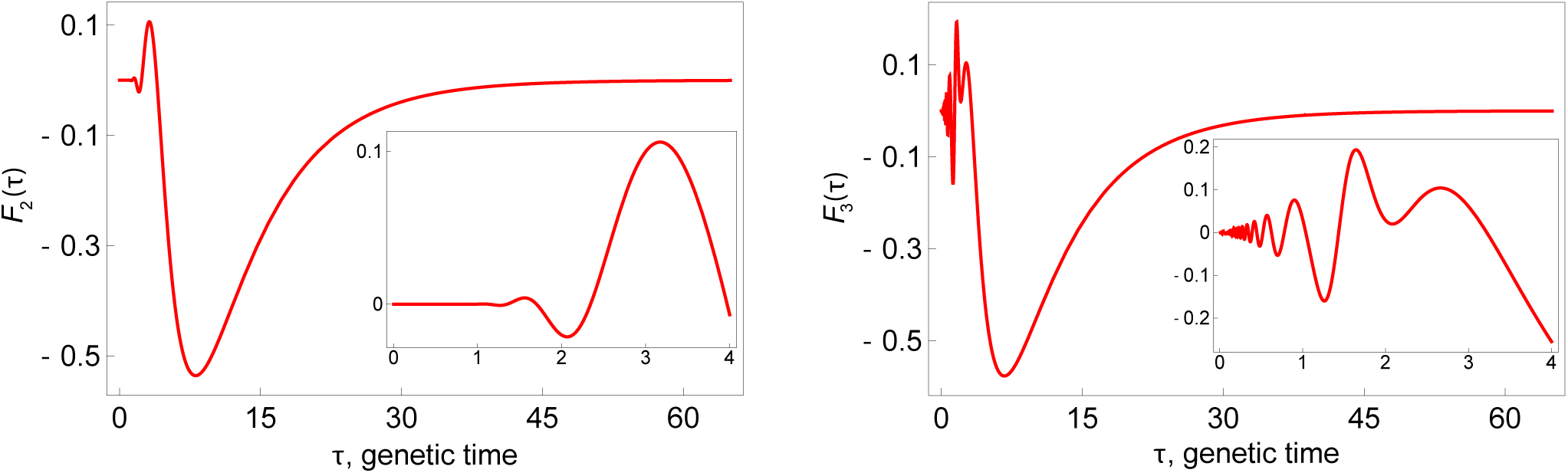
Two examples of a function which can be used to construct histories with identical frequency spectra. The present time is at *τ* = 0, whereas *τ* > 0 denotes the past. Left panel: The function *F*_2_(*τ*) constructed by Myers *et al.* to illustrate the non-identifiability problem. Right panel: The function *F*_3_(*τ*) discussed here. The insets show close-ups of the behavior of the corresponding functions near the present time.

However, Bhaskar and Song [25] pointed out that adding multiples of *F*_2_(*τ*) to any smooth history *Ñ*_1_(*τ*) leads to unrealistic population histories that oscillate increasingly rapidly as *τ* → 0^+^. In fact, they showed that any function satisfying Eq. (4) must exhibit an infinite number of sign changes. As such, any population size history function constructed by linear combinations of *F*(*τ*) is biologically unrealistic. They further proved that there is a unique solution to the inference problem in a very general class of realistic model functions. Thus, their argument offers the reassuring message that histories can, in fact, be uniquely reconstructed if we limit ourselves to biologically plausible histories and have sufficient data.

But how much data do we need? The oscillations near *τ* = 0 in *F*_2_(*τ*) are so small that they are barely noticeable in the inset within the left panel of Fig. 1 [for example, *F*_2_(*τ* = 0.5) ∼ 10^−12^]. It seems unlikely that these minuscule oscillations are relevant to our ability to reconstruct demographic histories. We expect (and show below) that replacing unrealistic small-amplitude oscillations by a realistic constant value would have an insignificant effect on the resulting spectrum. The resulting history is very different from a constant-sized history but would produce a nearly identical spectrum. If we formulate the inference problem in terms of model likelihoods, the Myers *et al.* construction shows that the likelihood surface is exactly flat along some directions in the space of all histories. Bhaskar and Song show that such flat directions are not present in the space of “reasonable” histories. We show below that the Myers *et al.* construction indicates the existence of almost-flat directions in the space of reasonable histories. For this reason, we cannot practically reconstruct history from the finite frequency spectrum alone.

The existence of almost-flat likelihood directions is not particularly surprising. Very ancient events are expected to leave few traces in the present-day allele frequency spectrum, and we cannot hope to reconstruct them from finite genetic data. In fact, the expansions and bottlenecks in the construction of Myers *et al.* (which are visible in the left panel of Fig. 1 and on Fig. 3) are ancient enough (tens of thousands of generations ago in a population of size 10,000) that they would have no statistically discernible individual effect on the present spectrum (see Appendix C).

**Figure 3:**
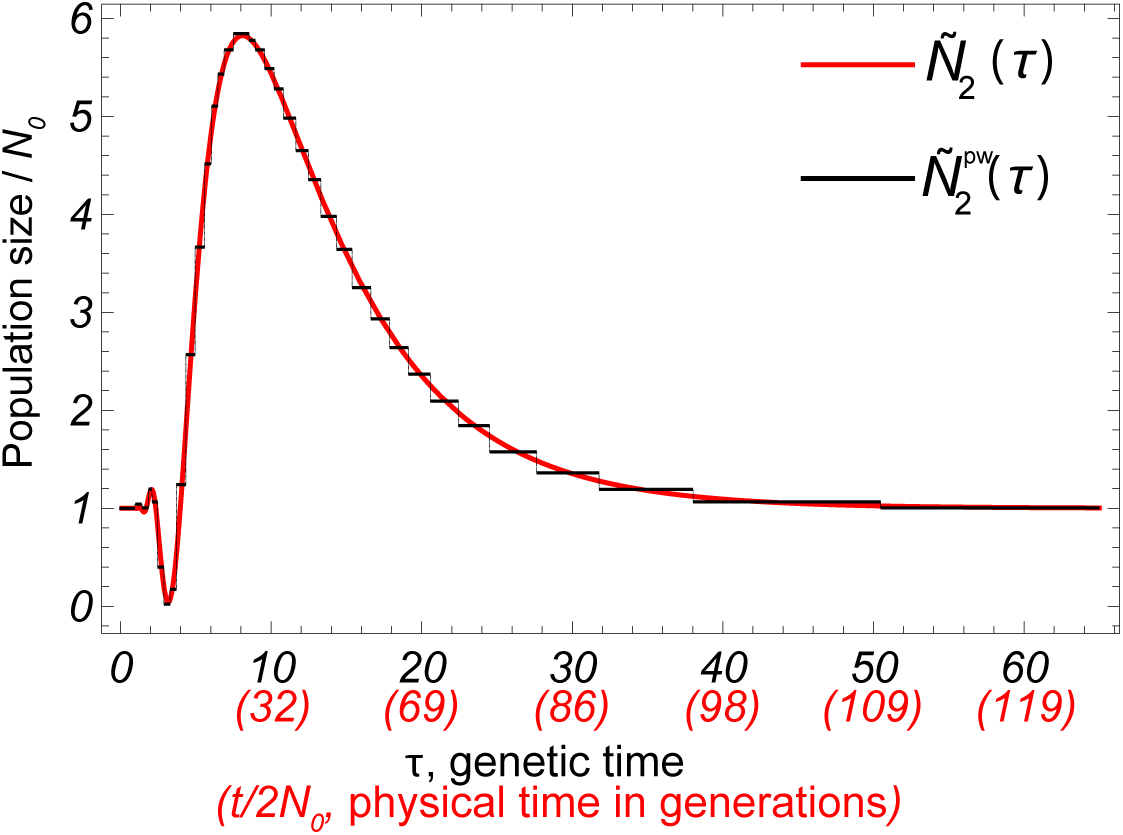
Population history *Ñ*_2_(*τ*) derived by Myers *et al.* (red, smooth) and its piecewise-constant approximation 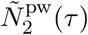 (black, piecewise-defined) with *p* = 35 pieces. Physical times corresponding to the indicated genetic times are presented in units of 2*N*_0_ generations. The time axis is not linear in physical time.

To investigate whether the Myers *et al.* construction can provide us with ‘unexpected’ flat directions in the likelihood surface, we consider another instance of *F*(*τ*) with larger-amplitude oscillations near *τ* = 0 and a more recent strong bottleneck, while still satisfying the orthogonality condition given in Eq. (4). We construct this function by following the prescription of Myers *et al.* [24] and performing the convolution of *f*_1_(*τ*) given above with 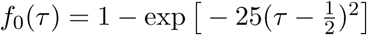, i.e.,

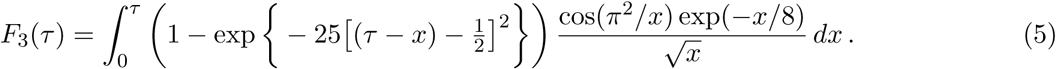

The function *F*_3_, depicted in the right panel of Fig. 1, has more prominent oscillations closer to the present time (*τ* = 0) compared to *F*_2_(*τ*) constructed by Myers *et al.*, as can be seen in the two insets. For example, *F*_3_(*τ* = 0.5) ∼ 10^−2^ whereas *F*_2_(*τ* = 0.5) ∼ 10^−12^.

To investigate the measurable effect of the fine-scale oscillations on the expected frequency spectrum, we first consider the piecewise-continuous function

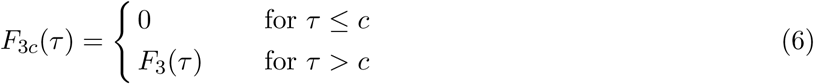

where we have introduced a cutoff *c* < 1 to remove the high-frequency oscillations near present in *F*(*τ*) given by Eq. (5). We numerically study the effect of this cutoff on the allele frequency spectrum. We use ∂a∂i [3] to compare the simulated expected allele frequency spectra of two demographies, one given by a constant population size

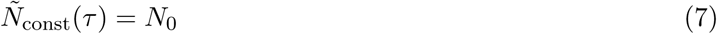

and one constructed using the truncated function defined in Eq. (6),

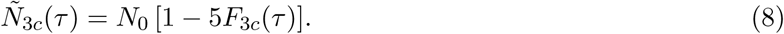

By construction, histories *Ñ*_const_(*τ*) and

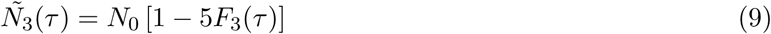

produce identical expected frequency spectra. Moreover, histories *Ñ*_3_(*τ*) and *Ñ*_3*c*_(*τ*) differ only for 0 ≤ *τ* ≤ *c*, the region dominated by the infinitely fast but small-amplitude oscillations. Given the small difference between these two population size histories, we expect the difference between the frequency spectra to be small as well. Fig. 2 shows simulated differences for different values of cutoff *c*. The left panel shows the expected allele frequency spectrum, highlighting the fact that different values of *c* produce practically indistinguishable frequency spectra, whereas the right panel depicts the relative change in the expected frequency spectrum as a result of the introduction of the cutoff threshold. As expected, decreasing the value of c lead to smaller relative errors.

**Figure 2:**
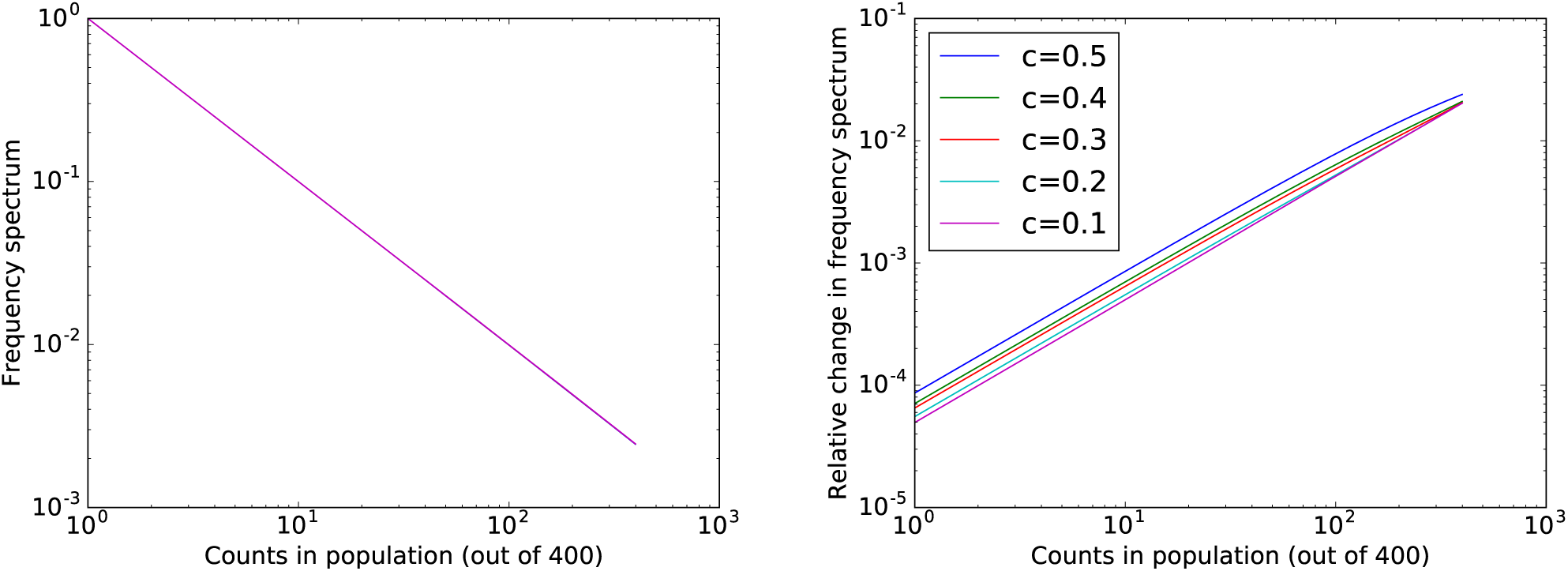
The effect of the cutoff c on the allele frequency spectrum, for *c* ∈ {0.5, 0.4, 0.3, 0.2, 0.1}. Simulations are done with ∂a∂i for a sampled population of size 400, and the plots are in log-log scale. Left panel: Practically identical allele frequency spectra of population histories *Ñ_c_*(*τ*) given by Eq. (8) for different values of c shown above. Right panel: Relative difference in the frequency spectra of *Ñ_c_*(*τ*) and *Ñ*_const_(*τ*) (that is, |*P_c_* – *P_const_*| /*P*_const_) for different values of *c*.

By eliminating the highly oscillatory part of the demographic model introduced in Ref. [24], one therefore arrives at a history which is biologically plausible and which leads to an allele frequency spectrum that is almost indistinguishable from that of the original history. To investigate whether such small differences could nevertheless allow for model identification given a large enough sample size, we next obtain analytical upper bounds on the difference in likelihoods between histories that are close enough to each other. Using this bound, we generalize the process described above to construct biologically realistic but very different histories that produce practically indistinguishable histories.

Given a realistic demographic history *Ñ*_1_(*τ*), we use the prescription of Myers *et al.* to generate a new history *Ñ*_3_(*τ*) whose allele frequency spectrum is identical to that of *Ñ*_1_(*τ*). We then construct a biologically realistic history *Ñ*_3*c*_(*τ*) such that |*Ñ*_3*c*_(*τ*) – *Ñ*_3*c*_(*τ*)| < ∈ *Ñ*_3*c*_(*τ*) for all *τ* and a small ∈. Finally, we use the bounds derived below to guarantee that *Ñ*_3*c*_(*τ*) is practically indistinguishable from *Ñ*_1_(*τ*) for arbitrary sample sizes.

## 3 Solving the diffusion equation

Before deriving the bounds discussed above, we first provide theoretical background and an overview of the derivation of Myers *et al.* in Ref. [24] which will form the basis of our calculations.

To solve the diffusion equation, we first consider *K*(*y,τ*_1_|*x*,*τ*_0_), the probability density that an allele whose frequency was *x* at a time *τ*_0_ has frequency *y* at a later time *τ*_1_. In other words, *K*(*y,τ*_1_|*x*,*τ*_0_) is the Green’s function for Eq. (2) and, therefore, satisfies

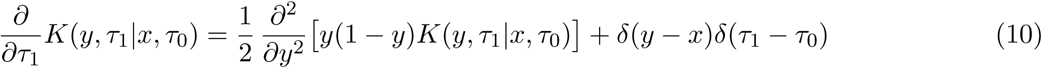

for 0 < *x*, *y* < 1. The solution to this equation is given by [26, 24]

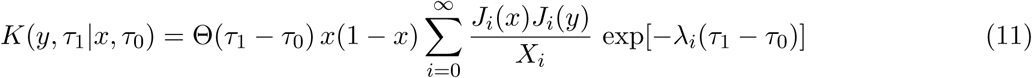

where Θ(·) is the Heaviside step function, λ_*i*_ = (*i* + 1) (*i* + 2)/2, and *J_i_*(·) are expressed in terms of the Jacobi polynomials as 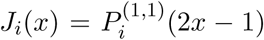 and are orthogonal according to 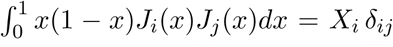 with *X_i_* = (*i* + 1)/[(*i* + 2)(2*i* + 3)]. Because *K*(*y*, *τ*_1_|*x*, *τ*_0_) is the Green’s function for Eq. (2), one can write the solution to Eq. (2) as

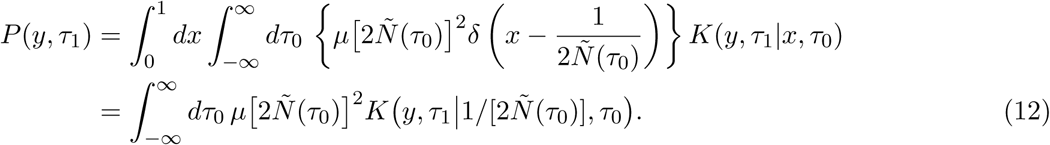

In other words, the present-day frequency spectrum can be obtained by integrating contributions from mutations that appeared at time *τ*_0_.

To proceed further, following Myers *et al.* [24], one can Taylor expand *K*(*y*, *τ*_1_|*x*, *τ*_0_) around *x* = 0 to find *K*(*y*, *τ*_1_|*x*, *τ*_0_) = *xQ*(*y*, *τ*_1_ - *τ*_0_) Θ(*τ*_1_ - *τ*_0_) + 𝒪(*x*^2^) for small *x* where 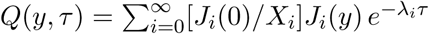. Therefore, assuming a large population size (i.e., min_*τ*0_[*Ñ*(*τ*_0_)] ≫ 1) leads to

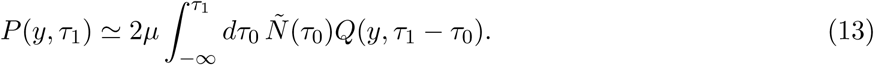

To simplify notation, we now measure time backwards from present (that is, *τ*_1_ =0 and to = −*τ*), define *P*(*y*) = *P*(*y*, *τ*_1_ = 0), consider the rescaled population size *ñ*(*τ*) = *Ñ*(−*τ*)/*N*_0_, and, without loss of generality, measure the frequency spectrum in units of 2_*μ*_*N*_0_. Hence, Eq. (13) becomes

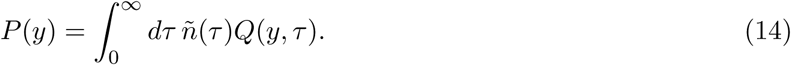

This is equivalent to Eq. (13) of Ref. [24], which defined the frequency spectrum up to an arbitrary constant. Using the expansion of *Q*(*y*, *τ*), one finds 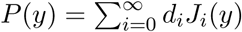 where

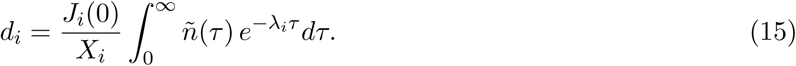

For example, a population with constant size *N*_0_ has *P*(*y*) = 2/*y* and *d_i_* = 2*J_i_*(0)(2*i* + 3)/(*i* + 1)^2^.

The function *F*_2_(*τ*) defined in Ref. [24] was designed to ensure that the coefficients {*d_i_*}, and therefore the frequency spectrum, are unaffected by the change in population history shown in (3). This condition leads to orthogonality condition (4) and the identifiability problem. In the next section, we investigate what happens when condition (4) is met only approximately, as happens when we construct a smoothed version of function *F*(*τ*).

## 4 Bounds on the change in the frequency spectrum and likelihood

We now bound the difference between two allele frequency spectra resulting from two different histories. Consider two histories, given by *ñ*′(*τ*) and *ñ*(*τ*), and their resulting frequency spectra, *P*′(*y*) and *P*(*y*) respectively. Using Eq. (14), we can write

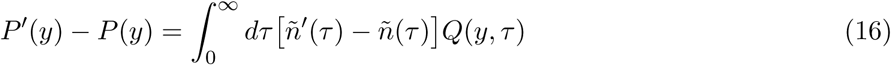

for the difference between the two allele frequency spectra. Let *δñ*(*τ*) = *ñ′*(*τ*) – *ñ*(*τ*) denote the difference between the two histories. Because *Q*(*y*, *τ*) ≥ 0, we find

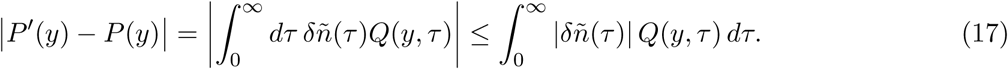

If there exists a small ∊ for which

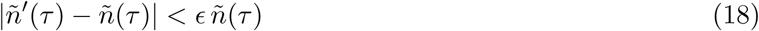

for all *τ* > 0, then

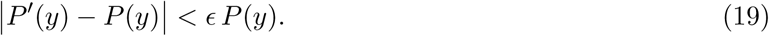

Therefore, ∊ also determines the upper bound on the difference between the two frequency spectra.

As an example, define 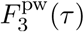 to be an approximation of the function *F*_3_(*τ*) by p piecewise-constant segments for *τ* > *c* and set 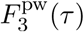 = 0 for *τ* ≤ *c* where *c* is an arbitrary, small number. Then consider the population size history given by the function 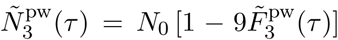. This approximates the original model of Myers, and the approximation can be made arbitrarily accurate by choosing a large but finite number of pieces *et al.* (see Fig. 3). Equation (19) tells us that the allele frequency spectrum from the approximate, piecewise-constant history can be made arbitrarily close to the history constructed by Myers *et al.* and, therefore, and thus also close to that of a history with constant population size. In short, we have shown that two biologically reasonable but *very distinct* histories *Ñ*_const_(*τ*) = *N*_0_ and 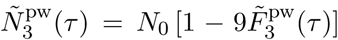, which belong to the same piecewise-constant family of functions and whose difference has a finite number of sign changes, lead to very similar allele frequency spectra. Our ability to detect the differences in the two spectra will therefore depend on the sample size and the length of the genome.

For a finite sample of size *M* and an infinite genome, the expected allele frequency spectrum becomes

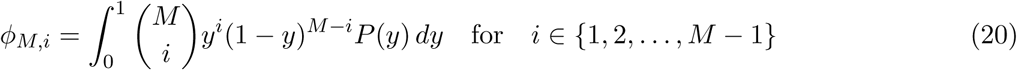

with *ϕ_M,i_* representing the number of loci where *i* out of *M* chromosomes in the sample have a derived allele. For a finite genome with *S* unlinked polymorphic loci in the sample of size *M*, we can compute a likelihood by assuming that allele frequencies are distributed according to the Poisson distribution [27, 3]. The likelihood 𝓛({*ϕ_M,i_*}) of the frequency spectrum (and of the history that generated it) equals the probability distribution for the observed allele frequencies spectrum {*m_i_*}:

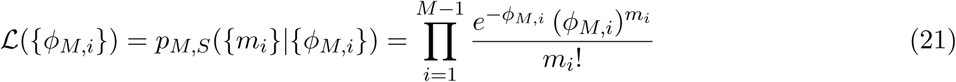

where 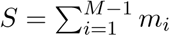 is the number of segregating sites.

Armed with this result, we can compare the likelihoods of two models originating from two very close population size histories.

First, we calculate a bound on the change in the expected finite–sample allele frequency spectrum for any two histories that are similar according to condition (18). We write the change in the allele frequency spectrum as 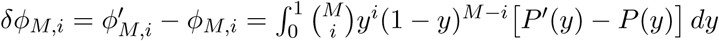. Using Eq. (19), we find

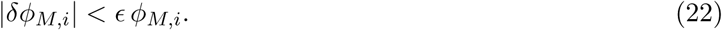

Equations (22) and (19) provide tight bounds, since there exist two population size histories that fulfil these bounds, namely *ñ′*(*τ*) = (1 + ∈) *ñ*(*τ*) and *ñ′*(*τ*) = (1 — ∈) *ñ*(*τ*). These bounds follow from the fact that finite and infinite-sample frequency spectra can be computed as integrals of the population history function with a positive kernel.

To compute the bound in likelihood differences between two models, we want to use the fact that one of the spectra *ϕ_M_,_i_* is optimal, in the sense that it is expected according to the maximum-likelihood history *ñ*(*τ*). Technically, we further require that *ñ*(*τ*) is not on the boundary of allowed histories, in the sense that, for γ small enough, *ñ*(*τ*) ± *γ*(*ñ′*(*τ*) – *ñ*(*τ*)) are acceptable histories, and thus with a likelihood worse than that of *ñ*(*τ*).

Under these conditions, Appendix A shows that the difference in likelihood obeys the tight bound

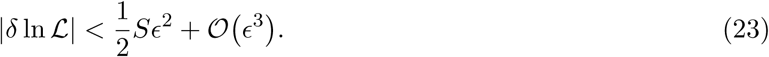

The same result holds if we use a multinomial distribution for allele frequencies rather than a Poisson random field (see Appendix B).

Because *ñ*(*τ*) maximizes the likelihood, the quadratic form of the bound was expected. However, the prefactor allows for specific predictions. Suppose that the maximum-likelihood estimate is the constant population size *Ñ*_const_(*τ*) = *N*_0_ (i.e., *ñ*(*τ*) = 1). We ask whether this can be distinguished from the history *Ñ*_3*c*_(*τ*) obtained by removing the recent fast oscillations from *Ñ*_3_(*τ*) up to *c* = 0.028556. The expected number of segregating sites for a sample of size *M* according to the constant history is *S* ≃ 4_*μ*_*N*_0_ ln *M*.

The difference in population size history provided induced by the flattening corresponds to ∈ = 10^−4^.

To determine whether *Ñ*_const_(*τ*) and *Ñ*_3*c*_(*τ*) are distinguishable, we consider |*δ* ln 𝓛| ∼ 2 as the minimal difference in likelihood to enable reliable distinction between the two models. Assuming a uniform prior on the constant and variable models, 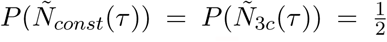, a Bayesian estimate for model posteriors is bounded as 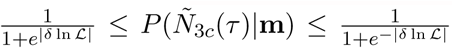. In other words, |*δ* ln 𝓛| < 2 means that 0.12 ≤ *P*(*N*_3*c*_(*τ*)|**m**) ≤ 0.88: If *Ñ*(*τ*) = *N*_0_ is the maximum likelihood estimate, we cannot rule out *Ñ*(*τ*) = *Ñ*_3*c*_(*τ*).

Taking |*δ* ln 𝓛| ∼ 2 as the measure of a detectable change in the log-likelihood and using a genome-wide mutation rate of *μ* ≃ (1.4 × 10^−8^ bp^−1^ gen^−1^) × (3.6 × 10^9^ bp) = 50 gen^−1^ [11], we find that for *N*_0_ = 10,000, the sample size *M* needed to distinguish these two models is approximately 10^43^, which is much larger than the current human population size. Similarly, given a sample of size of the entire population of 10,000 and a per-locus mutation rate of 1.4 × 10^−8^ bp^−1^ gen^−1^, we would need a genome with length *L* > 3.8 × 10^10^ to identify the two spectra. This is longer than the human genome. There simply is not enough data in the human genome to allow us to resolve the difference between the two models, leaving aside the fact that for such large sample sizes and small effects, failures of the diffusion approximation might have a much larger effect than the differences that we are trying to detect between the population histories [28].

### 4.1 Comparison with the findings of Terhorst and Song

Terhorst and Song also proposes bounds on identifiability with finite genomes [6] and bounds on identifiability that are independent of sample size. Specifically, it shows that *any* estimator of the demographic history that is based on the frequency spectrum has an expected worst-case error proportional to 1/log(*S*), i.e., there exists a demographic history *ñ*(*τ*) for which the expected *τ_c_*-truncated error 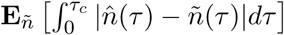 between the true history and inferred history 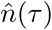 is proportional to 1/ log *S*. In a sense, this appears to be stricter and more pessimistic than our bound: for a given difference in likelihood, our detectable error threshold scales as 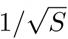, which decreases much faster in a long genome.

The bounds have quite different interpretation, however: we are asking about the best we can possibly do, whereas Ref. [6] considers the worst-case scenario. In other words, we show that *any* two histories within an error range are indistinguishable, whereas Ref. [6] shows that *there exist* histories which are indistinguishable for a given error threshold.

This is an important difference. The 1/log *S* bound relies on constructing a set of histories that change with S by assuming the existence of a historical bottleneck whose size e decreases as 1/log(*S*). Intuitively, a tighter bottleneck limits our ability to see through the bottleneck into previous history by reducing observable diversity. In fact, we can use the Terhorst and Song formalism to obtain even more conservative bounds by making the bottleneck size ∈ decrease faster than 1/log(*S*). At that rate, the amount of information lost to the narrower bottleneck will not be compensated for fast enough by the increasing *S*. Thus differences in histories prior to the bottleneck can be made practically unconstrained for a given *S*. In the Terhorst and Song notation, we could choose *δ* = (*M* – ∈)/*J* in their Equation 21, then make ∈ arbitrarily small (specifically, for an arbitrarily small constant *ν*, we can choose ∈ small enough such that 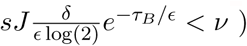, leading to a bound

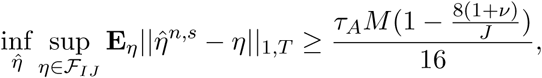

where *τ*, *M* and *J* are parameters defining the family of functions considered in Ref. [6]. Selecting *J* ≥ 9, and *ν* ≪ 1, this bound is positive and independent of *S*. It is also stricter than the 1/log(*S*) bound of Terhorst and Song. The 1/log *S* convergence of Ref. [6] is therefore correct, but a consequence of the particular choice of the scaling of the bottleneck size. In general, worst-case bound are dominated by unconstrained histories with very tight bottlenecks. It is not clear that these are the most relevant for practical inference.

Ref. [6] also presents, in Theorems 4 and 7, bounds given a fixed bottleneck size, which are more readily compared with the bounds presented here. In that case, the bound on the expected error decreases as 1/*S*. Thus this minimax bound from [6] is less strict for long genomes than the best-case bound presented here, while being valid for a much wider range of estimators. There are many differences in the specific families of functions and estimators and the bounding strategies presented here and in [6] that could explain this difference in convergence rate.

From a practical perspective, the bound obtained here does provide a simple, clear, and practical bound for likelihood-based estimation which we could use directly to estimate the possible effect of minor fluctuations in population histories.

## 5 Conclusion

Demographic histories inferred from genetic data help us understand human history, evolution, and the distribution of deleterious variants. For this reason, unacknowledged errors in inference can lead to flawed conclusions in many downstream interpretations. The work of Myers *et al.* highlighted an uncontrolled source of potential error and led to more cautious interpretation of demographic inference studies [25, 5, 6]. However, it did not provide a strategy to quantify or limit this uncertainty. Bhaskar and Song [25] suggests that the uncertainty may not be as large as suggested by Myers *et al.,* once we limit our attention to biologically realistic functions.

Even though the limitation to biologically realistic functions does resolve the identifiability problem in an idealized setting, we showed that it has little bearing on practical inference; because the genome is finite, we still do not have the statistical power to distinguish between vastly different histories. The work of Bhaskar and Song (and that of Terhorst and Song [6]) provides, in principle, a systematic way of assessing the uncertainty by considering very general families of possible histories. This is computationally demanding, however, and many studies continue to consider histories parameterized by a small number of parameters.

The main practical message from the present work is that is any inference that identifies a demographic history with high precision based on human data must rely on strong implicit or explicit assumptions, and is missing out smooth, biologically realistic alternatives. Low parameter uncertainties should be taken as a warning sign rather than a reassurance that uncertainties have been taken into account. To ensure that downstream analyses do not depend strongly on these uncertainties, validation over multiple histories consistent with the data is necessary.

## Acknowledgements

This work was supported by CIHR through the Canada Research Chair program and by a Sloan Research Fellowship (to S.G.).

## A Bounds on the change in log-likelihood: Poisson formulation

In this section, we derive the change in log-likelihood of a demographic model as a result of a change in the population size history. Given the allele frequency spectrum *P*(*y*), the expected allele frequency spectrum in a sample of *M* individuals is 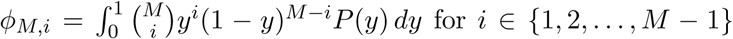.

In a finite genome with 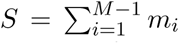 unlinked polymorphic loci in the sample of size *M*, we assume that the probability of observing frequency distribution {*m_i_*} is a product of Poisson distributions, i.e., 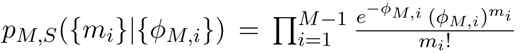 where 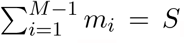, the number of segregating sites. We write the difference in log-likelihood between the two models as

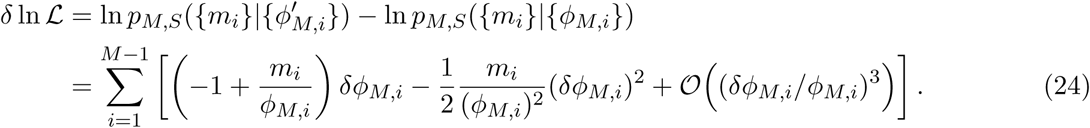

We now assume that *ϕ_M,i_* is the frequency spectrum obtained by maximizing the likelihood function L with respect to the model *ñ*(*τ*); in other words, we compute the change in likelihood for models close to the maximum likelihood model. We wish to use this condition to set the linear term in *δϕ_M,i_* to zero. To this end, first note that *ϕ_M,i_* is a linear functional of *ñ*(*τ*): we can relate finite changes *δñ*(*τ*) in the demographic model to changes {*δϕ_M,i_*}_*i*=1,…,*M*_ in the frequency spectrum as

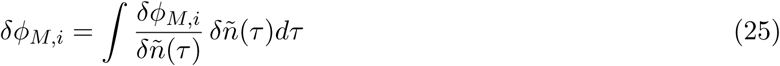

where the functional derivative

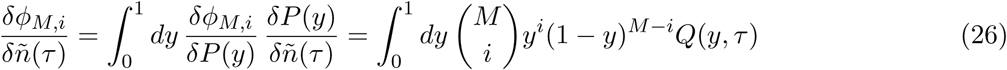

is independent of *δñ*(*τ*).

If the linear term 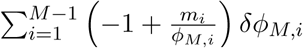 is not zero, we can construct a history *ñ*(*τ*) + *γδñ*(*τ*), for a small constant |*γ*| ≤ 1, with a better likelihood than *ñ*(*τ*). This is a contradiction, and the linear term must be zero, unless the history *ñ*(*τ*) + *γδñ*(*τ*) is not a valid demographic model. This can happen if *ñ*(*τ*) is constrained at the boundary of allowed histories, for example if *ñ*(*τ*_0_) = 0 for some *τ*_o_, or if the space of allowed *ñ*(*τ*) is nonlinear such that two histories can be close to each other without intermediate histories being allowed. Our result therefore requires that histories of the form *ñ*(*τ*) + *γδñ*(*τ*) are valid for *γ* sufficiently small.

Assuming that this condition is respected, we have 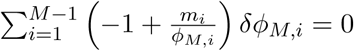 and we can write

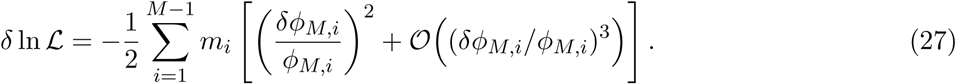

Therefore, we find

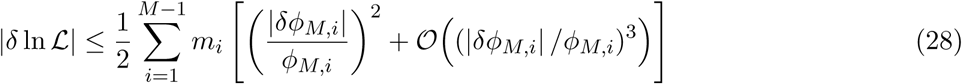

which, with Eq. (22) and the definition of *S*, becomes

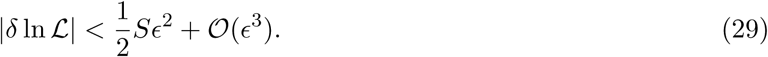

Here 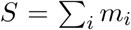 is an observed random variable and the result holds independently of the particular random realization of the *m_i_*. A similar bound can be obtained using Fisher Information:

## A.1 Fisher Information estimate of parameter variance

If we parameterize the demographic history as 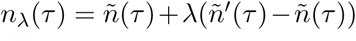, we can estimate the variance of parameter *λ* using Fisher’s information matrix

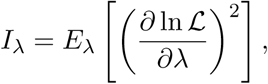

where the expectation is taken at fixed λ over the possible observations.

Computing derivatives as above, and using the assumption that the *m_i_* are independent with mean and variance *ϕ_M,i_*, we find

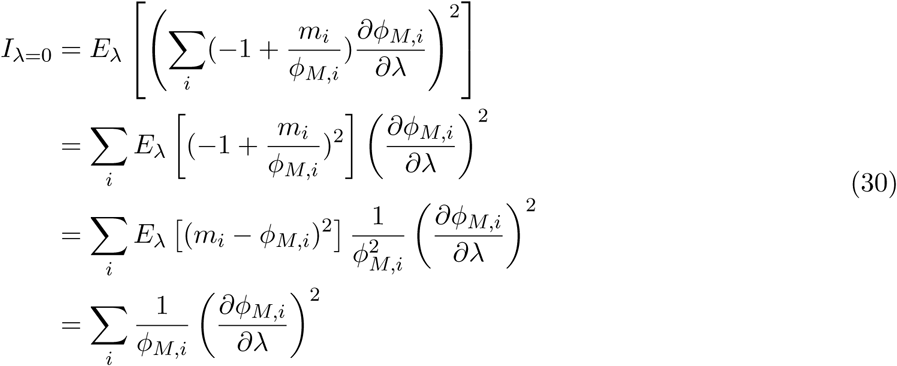

Since *ϕ* is a linear functional of *ñ*(*τ*) and a linear function of λ, we find 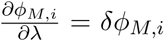, the difference in frequency spectra generated by histories *ñ′*(*τ*) and *ñ*. Thus

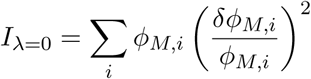

By virtue of Equation (22), we can bound this by

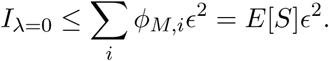

By Fisher’s theorem, if λ = 0, the MLE estimator 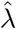 is asymptotically distributed with variance 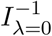. The bound on the log-likelihood discussed in the text roughly corresponds to asking that the standard deviation 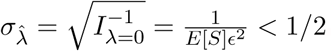, such that λ = 1 lies just outside a 95% confidence interval.

## B Bounds on the change in log-likelihood: Multinomial formulation

In this Appendix, we rederive the change in log-likelihood of the model as a result of a change in the population size history using a different distribution for the allele frequencies.

As discussed above, given the allele frequency spectrum *P*(*y*), the expected allele frequency spectrum in a sample of *M* individuals is 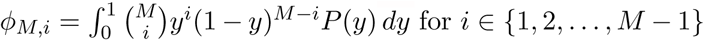. We now assume that in a finite genome with 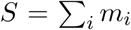 unlinked polymorphic loci in the sample of size *M*, the distribution of population frequencies is multinomial, i.e., 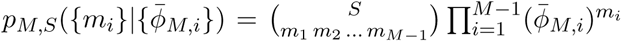 where 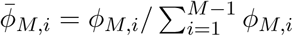 is the normalized allele frequency spectrum. In other words, each locus is drawn independently from the expected distribution 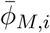. Hence, the log-likelihood difference between the two models becomes

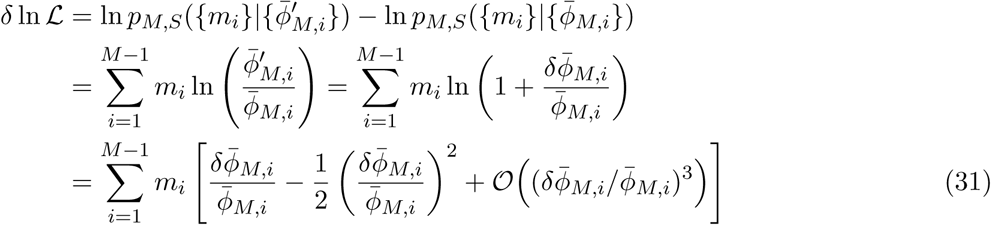

where, in the last step, we used the Taylor expansion ln(1 + *x*) ≃ *x* − *x*^2^/2 + 𝓞(*x*^3^) for *x* ≪ 1.

We now transform the functional dependence of *δ* ln 𝓛 from *δϕ_M,i_* (change in the normalized frequency spectrum) to *δϕ_M,i_* (change in the unnormalized frequency spectrum). Let *h* = *f/g* where *f*, *g*, and *h* are three functions. For small changes in *f* and *g*, we can write the corresponding change in *h* as

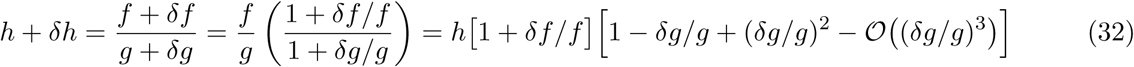

where we have Taylor expanded the denominator up to second order. We find

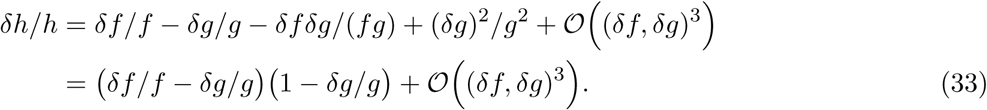

Now, let us substitute *f* → *ϕ_M,i_* and 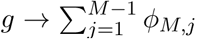 (that is, 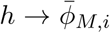). Therefore, the relative change in normalized frequency, up to second order, becomes

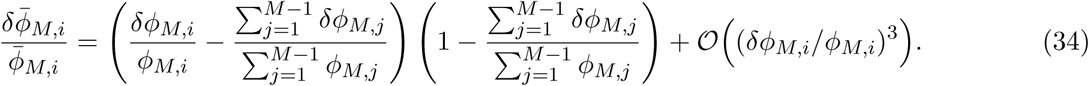

Substituting Eq. (34) into Eq. (31) leads to the series expansion of *δ* ln 𝓛 in powers of *δϕ_M,i_*. We formally represent this expansion as *δ* ln 𝓛 = *δ*^(1)^ ln 𝓛 + *δ*^(2)^ ln 𝓛 + 𝓞(*δϕ_M,i_*/*ϕ_M,i_*)^3^)

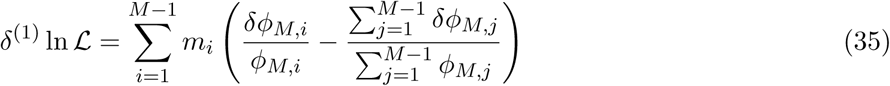

where

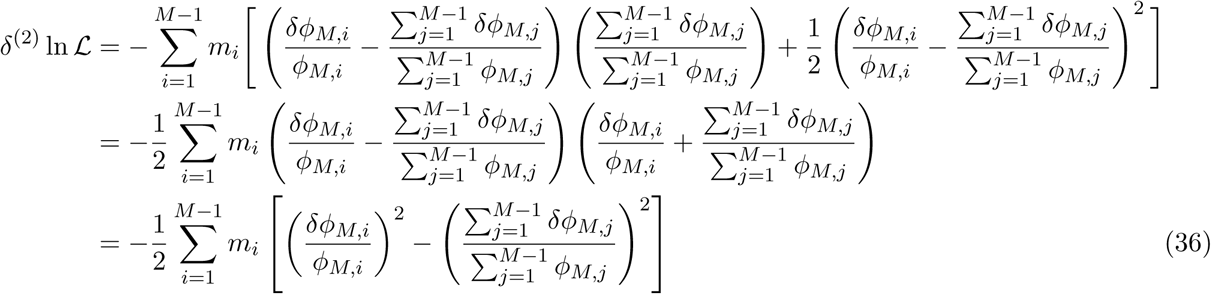

are, respectively, the first- and second-order changes in the log-likelihood expressed in terms of changes in the unnormalized frequency spectrum.

Next, we relate the relative change in the normalized frequency spectrum, given by Eq. (34), to a change in the demographic model. As shown in Appendix A, the change in frequency spectrum *δϕ_M,i_* depends only linearly on the change in the demographic model *δñ*(*τ*). This fact allows us to use Eqs. (31), (34), and (25) to derive the change in the log-likelihood function in terms of the change in the demographic model. In other words, *δ*^(1)^ ln 𝓛 and *δ*^(2)^ ln 𝓛 also represent respectively the first- and second-order changes in the log-likelihood as a result of a small change in the demographic model *δñ*(*τ*).

We now assume, as we did in Appendix A, that *ϕ_M,i_* is the frequency spectrum obtained by maximizing the likelihood function 𝓛 with respect to the model *ñ*(*τ*). Therefore, the first-order change in the log-likelihood function *δ*^(1)^ ln 𝓛 due to the change *δñ*(*τ*) around the best-fit model should vanish at the extremum, that is, *δ*^(1)^ ln 𝓛 = 0, and using Eq. (36), we can write

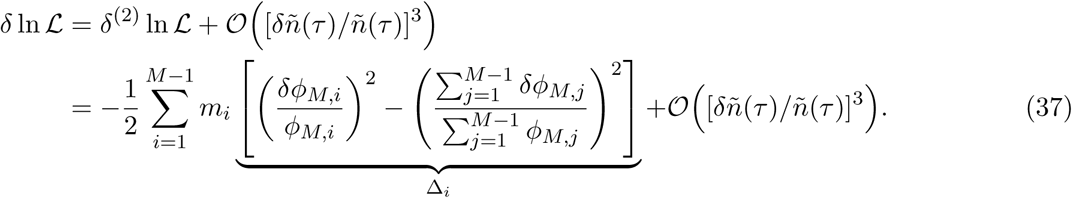

To derive the bounds on the change in log-likelihood, we now find the following bounds on Δ_*i*_ (defined in the previous equation)

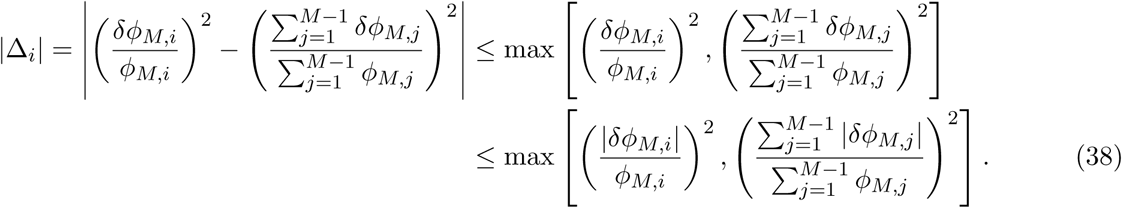

Using Eq. (22), both terms on the right-hand side are smaller than ∈^2^, so we can write

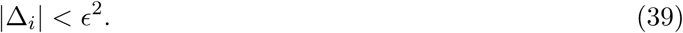

Finally, substituting (39) in Eq. (37) leads to the bound on the change in log-likelihood function

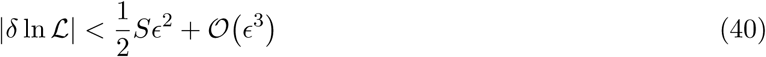

which is identical to that derived using the Poisson distribution for the frequencies.

## C Effect of bottlenecks

Here, we investigate the effect of the existence of a bottleneck in the demography, e.g., one given by (8), on the resulting allele frequency spectrum. Our motivation is to determine whether a bottleneck would leave a measurable impact on the present-day allele frequency spectrum given the timing (recent or past) of the bottleneck. In Fig. 4, we show (black, solid line) a demographic history constructed from the history given in Eq. (8), with the cutoff *c* = 0.5, by removing all bottlenecks; the original history is shown in red, dashed line. The resulting frequency spectrum (obtained using ∂a∂i), shown in Fig. 5, indicates that removing the recent bottlenecks (which are more recent compared to those in the example presented in Myers *et al.* [24]) indeed leaves a detectable effect on the spectrum, especially for the common variants. Performing the same analysis using the example given by Myers *et al.* [24] leads to negligible difference in the frequency spectrum (results not shown for brevity), mainly due to the fact that the bottlenecks are in distant past and have no discernible effect on the frequency spectrum observed at present.

**Figure 4:**
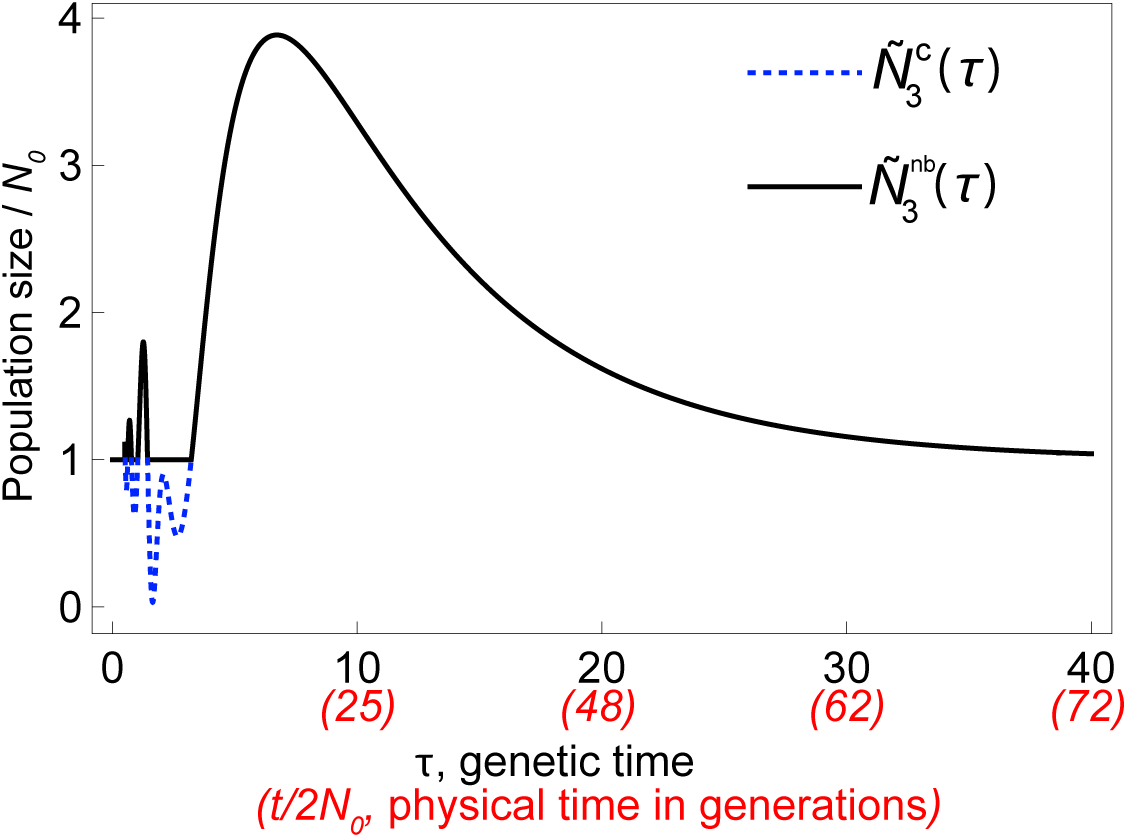
A population size history 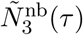 derived by removing bottlenecks from *Ñ*_3*c*_(*τ*) for *c* = 0.5 with the bottlenecks removed. Removed bottlenecks in the original history are plotted in blue (dotted lines). Physical times corresponding to the indicated genetic times for *Ñ*_3*c*_(*τ*) are presented in units of 2*N*_0_ generations. The time axis is not linear in physical time.

**Figure 5:**
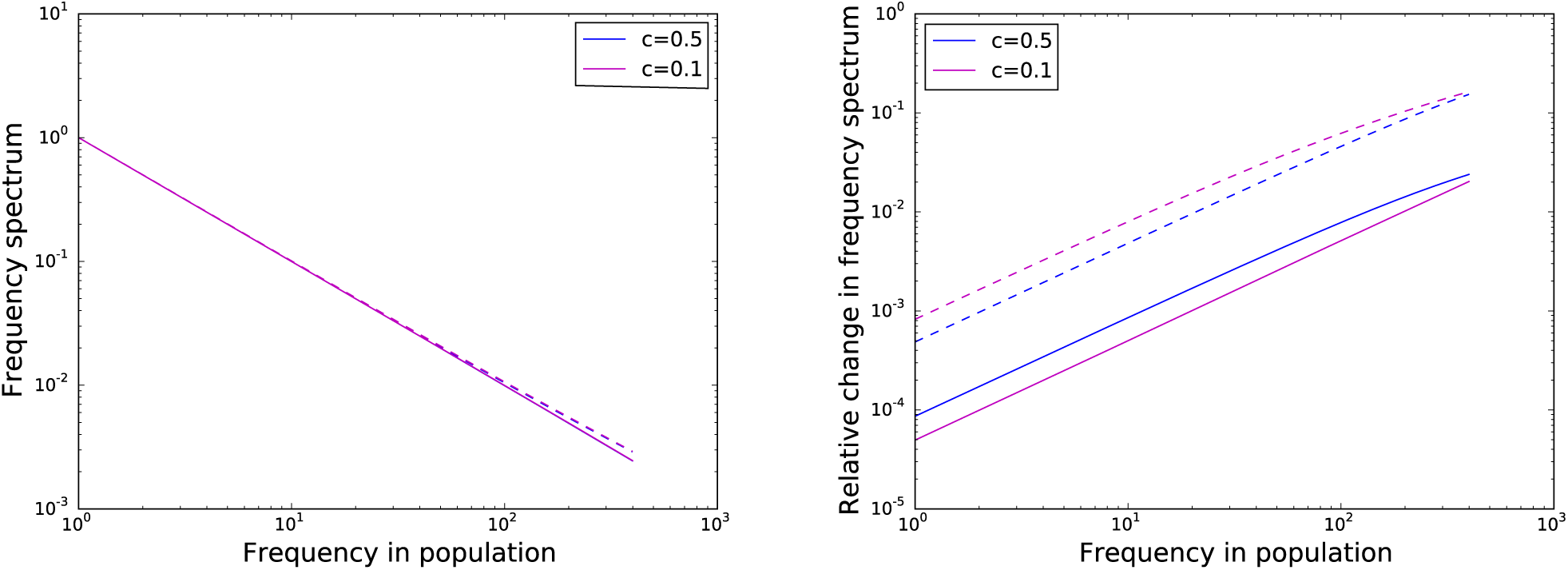
The effect of the removal of bottlenecks on the allele frequency spectra for different values of the cutoff c in two scenarios: (I) using the original history *Ñ*_*c*_(*τ*) defined in Eq. (8) (solid lines), and (II) using the aforementioned history without the bottlenecks, as discussed in Fig. 4 (dashed lines). Simulations are performed with ∂a∂i for a sampled population of size 400. Left panel: Removing bottlenecks causes an observable change in the frequency spectrum, especially for more common variants. Note that in both cases (that is, with or without bottlenecks) the effect of the small cutoff c on the spectrum is practically negligible. Right panel: Relative change in frequency spectrum (measured with respect to that of a population of constant size).

